# Top-down Proteomics of 10,000 Single Brain Cells

**DOI:** 10.1101/2023.05.31.543176

**Authors:** Pei Su, Michael A. R. Hollas, Stanislav Rubakhin, Fatma Ayaloglu Butun, Joseph B. Greer, Bryan P. Early, Ryan T. Fellers, Michael A. Caldwell, Jonathan V. Sweedler, Jared O. Kafader, Neil L. Kelleher

## Abstract

We introduce single-cell Proteoform imaging Mass Spectrometry (*scPiMS*), which realizes the benefit of direct analysis of intact proteins to process 10,836 single cells from the rat hippocampus. This new platform addresses the throughput bottleneck for single cell proteomics using mass spectrometry, boosting cell processing rates by >20-fold in the field. We identified 169 proteoforms <70 kDa from single brain cells and classified 2758 of them as neurons, astrocytes or microglia cell types.

## Introduction

Single-cell proteome analysis (SCPA) seeks to capture the heterogeneity in cellular type, state and function obscured by population-averaged measurements of proteins. Proteins in single cells and subcellular structures can been mapped using different molecular probe-based methods including antibody-based optical microscopy at high spatial resolution and sensitivity. However, such approaches require prior knowledge of target proteins and lack the molecular specificity to target proteoforms,^1^ limiting our ability to discover the exact forms of proteins present in different cell types. Therefore, a more precise characterization of normal and pathological states of cells comprising cellular neighborhoods in complex tissue currently evades us.^2^

Combining the specificity of proteoform analysis with single cell measurement presents major challenges. Unlike genomics platforms for single-cell RNA sequencing (scRNA-seq),^3-5^ single-cell proteomics (SCP) relies upon an analyte detection paradigm without amplification to interrogate the complexity of the proteome.^6^ Since 2018, mass spectrometry (MS) has been advanced to perform SCPA, using ‘bottom-up’ platforms such as Single Cell ProtEomics by Mass Spectrometry (SCoPE-MS).^7-11^ This work has increased the number of proteins that can identified and quantified using peptide-based workflows,^12^ yet the number of cells processed in a given study is limited to a few hundred per study.^13^ To increase throughput, avoid multi-step processing^14^ and protein digestion, a more direct sampling approach has the theoretical advantage of characterizing cells at the level of whole proteoforms. However, a scheme for intact proteoform SCPA using ‘top-down’ MS^15^ sets requirements for sensitivity and protein identification that have yet to be met.

Since the late 1990’s mass spectrometry has been capable of detecting individual ions.^16, 17^ However, even with the very recent incarnation of individual ion mass assignment^18^ the question of how single particle MS could drive single cell analysis has been left open. A suitable top-down workflow should both detect and identify whole proteins for SCP.^19^ This could also intersect with MS imaging (MSI), which involves direct sampling of small amount of highly localized protein(s).^20^ Importantly, the use of an imaging mode would empower fast analyses to identify even rare cells from a mixture of thousands. In 2022, we developed proteoform imaging mass spectrometry (PiMS),^21^ which broke through some long-standing challenges by detecting and identifying hundreds of proteoforms up to ∼70 kDa from thin tissue sections at ∼50 μm spatial resolution.^21^ The PiMS platform utilizes a moving liquid bridge to generate proteoform ions by electrospray ionization and proteoform readout is accomplished by individual ion mass spectrometry (I^2^MS),^22, 23^ a new technique for single ion detection with >500x greater sensitivity and 10x higher resolving power over ensemble MS.^24, 25^ Thus, the new capabilities of PiMS have enabled fast and direct analysis of attomole proteins amounts.

Here, we introduce **single-cell Proteoform imaging Mass Spectrometry (*scPiMS*)** to enable facile proteoform analysis from single cells spread onto glass slides.^26^ *scPiMS* profiling of 10,836 cells disaggregated from rat hippocampus was performed in 10 days, yielding detection of 472 single-cell proteoforms with over 160 identified including well-known markers like ENOG (neurons) and GFAP isoforms (astrocytes) that striate brain cell types without antibody staining. *scPiMS* demonstrates a high potential to advance single cell biology analogous to the early days of scRNA-seq.^27^ Utilizing selected cell-type markers, *scPiMS* was able to stratify 2758 individual cells into three different cell types; astrocytes (1538), microglia (712), and neurons (508).

## Results and Discussion

### The single-cell PiMS workflow

A group of a few thousand cells were disaggregated from a rat hippocampus and drop cast onto a glass slide such that their average spacing was on the order of ∼200 μm (**Fig. 1a**, far left).^26^ To directly sample proteins from these cells, we employed proteoform imaging^21^ to raster a ∼100 μm liquid droplet across the glass slide (**Fig. 1a, middle**) that is continuously sampled by an electrospray ionization source. As a result, when the droplet reached a single cell (**Fig. 1a**, middle, image II), thousands of proteoform ions were detected by individual ion mass spectrometry.^24^ This burst of ions from a single cell was plotted as an extraction curve termed a “cellogram” (**Fig. 1a**, right), and the 8833 individual proteoform ions collected over six seconds are shown in **Fig. 1b**.

**Fig. 1.**
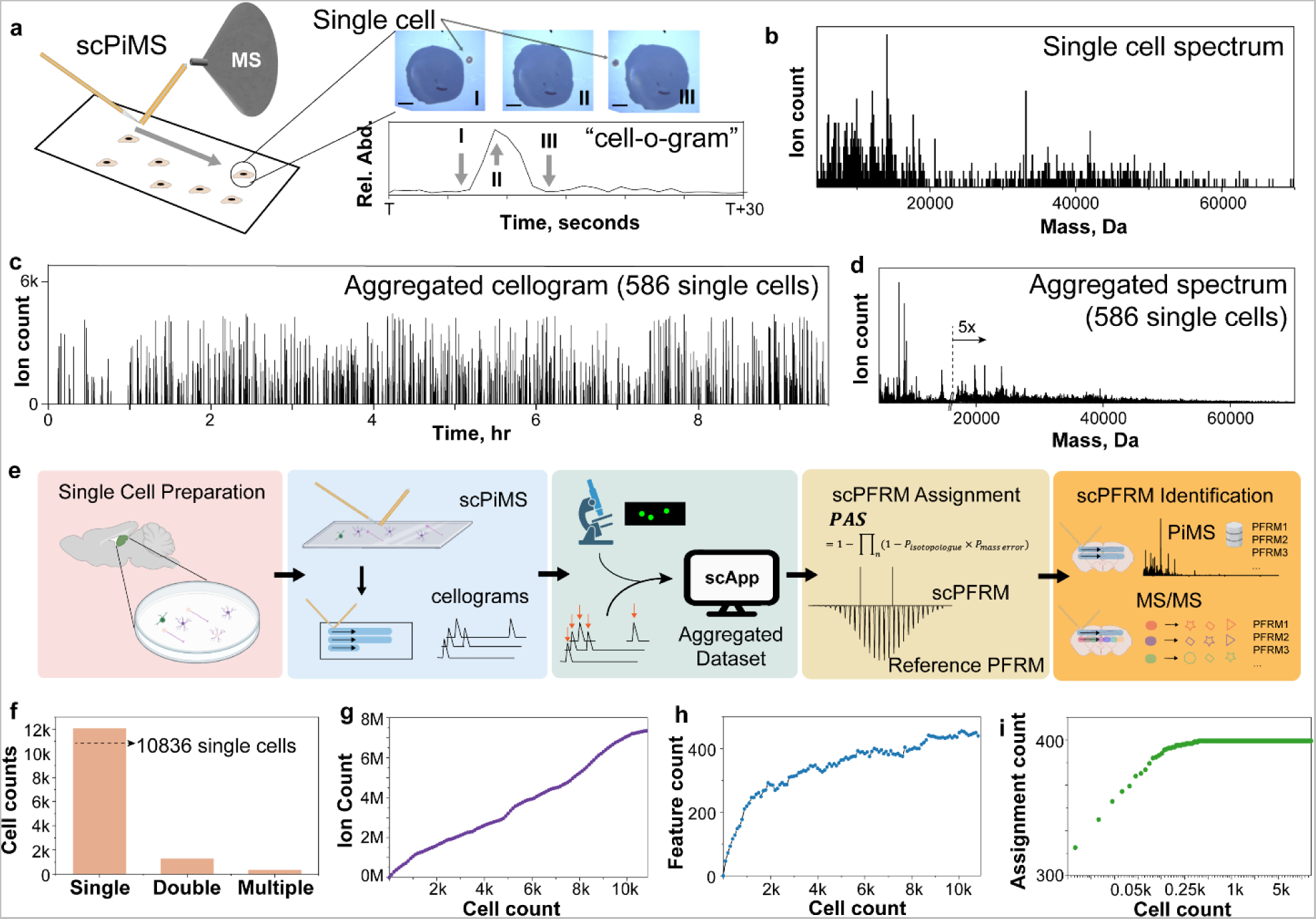
The *scPiMS* workflow for proteoform detection and identification from single cells. Panel (a) shows the scanning approach (at left); snapshots at right show the relative location of a cell before (I), on (II), and after (III) analyte extraction by a liquid bridge (scale bar = 50 µm). The extraction profile of the single cell (a “cellogram”) is shown at right. Mass spectrum of the ions collected from a single cell is shown in panel (b). The aggregated cellogram containing 586 single cell features in one run is shown in (c), and its corresponding aggregated mass spectrum is shown in (d). (e) Workflow for the main steps involved with the *scPiMS* process. (f) Ratio of single cell versus double and multiple cell features in the *scPiMS* run yielding the 10.8k molecular features data set. Collector’s curves for number of charge-assigned ions (g), proteoforms detected algorithmically as well-formed isotopic distributions (h), and proteoforms assigned to the rat hippocampal ion library using PAS (i).

To expand the number of cells sampled, we used *scPiMS* at a probe rastering rate of 15 μm/s to scan a slide similar to a PiMS tissue imaging experiment.^21^ The scan rate was set such that the liquid bridge leaves a cell by the time protein extraction wanes (Methods). For a 0.5 cm^2^ region of a slide sampled over ∼10 hours, we obtained 586 single cell extraction events with the aggregated cellogram from this run shown in **Fig. 1c** (Methods). During this acquisition period, 0.72 million individual ions were measured and aggregated (**Fig. 1d** and Methods). In the aggregated mass spectrum, we used an intact mass tag (IMT) approach to manually identify a few of the most abundant proteoforms by checking against a list of reported rat hippocampal proteoforms within a ±1.5 part-per-million (ppm) mass tolerance (Methods).^28, 29^ The most abundant ones in the aggregated single cell spectrum were isoforms of Myelin basic protein (MBP), High mobility group nucleosome-binding domain 2 (HMGN2), Histones H1 and H2A, Thymosin beta-4 (TYB4), Thymosin beta-10 (TYB10), and ATP synthase subunits.^28, 29^

### The scPiMS 10.8k brain cell dataset

We next used an implementation of *scPiMS* with ∼3-fold higher throughput to examine proteoforms and their distributions across several slides containing a total of ∼15k cells. This experiment took 200 hours of unattended acquisition time using a 200 µm probe at a rastering rate of 40 µm/s (*i.e.*, a nominal processing rate of 80 cells/hour). Using the full workflow for *scPiMS* depicted in Fig. 1e, we detected 12062 cellogram peaks and used an algorithm to extract 10,836 single cell detection events (Methods). This demonstrated a 90% success rate in converting microscopy-registered features (centered at cell nuclei stained with DAPI) into single cell proteoform detection (**Fig. 1f**). Upon spot checking 200 randomly selected cellular detection events (video recorded at 900x magnification during the *scPiMS* process), all were deemed to be true single cell sampling events.

### Assignment of scProteoforms

The 10.8k dataset was comprised of 48,412 intact ion scans capturing a total of ∼16 million raw ions. To assist with assignment of integer charge states to these ions, the dataset was processed by I^2^MS software^18^ after combination with a reference ion library obtained from PiMS imaging of rat hippocampal tissue. The entire dataset for the 10.8k cells was aggregated together for improved assignment of mass to proteoform isotopic distributions and totaled 7.17 million charge- and mass-assigned individual ions from 5-70 kDa. Fig. 1g shows the rate of accrual for mass-assigned proteoform ions as data were aggregated to 10.8k cells. The aggregated *scPiMS* spectrum yielded 472 proteoform features matched to the ion library with a collector’s curve for them shown in Fig. 1h. Following this, an application was used to extract ions from single cells (“scApp”, middle panel of **Fig. 1e**). With thousands of ion masses detected per cell, we developed a statistical framework to assign proteoforms detected in each single cell mass spectrum using a proteoform assignment score (PAS, Methods). This framework and new score assigns a probability that a proteoform is present in a single cell and provides an estimated false discovery rate for this assignment (FDR, see the 4^th^ panel from the left in **Fig. 1e** and Methods). Using this approach, we obtained an average of 106 scProteoforms detected per cell above 5% estimated FDR.

### Identification of scProteoforms

Proteoform identification employed a two-pronged approach (**Fig. 1e**, far right). We first found 169 putative proteoform IDs using an intact mass tag (IMT) search against ∼5,000 candidate proteoforms derived from a group of rat protein sequences transformed from a mouse-hippocampus bottom-up proteomics study using ±1.5 ppm tolerance.^30^ Next, we obtained another set of experimentally-verified proteoforms obtained using MS/MS mode during PiMS^21^ on brain tissue sections from the same three parent animals in this study (**Table 1**). From this two-pronged process, we generated a total of 169 identified scProteoforms, with 18 of these MS/MS confirmed (**Table 1**). The top-three GO terms show enrichment of proteoform detection mapping to key proteins of central metabolism with *p-values* ranging from 10^-11^ to 10^-23^. The rate of accrual for already-assigned proteoforms is shown in Fig. 1i. We note that the rate of rise for this collector’s curve for proteoform assignment rises far more sharply vs. that for *de novo* collection of new isotopic distributions (*i.e.*, compare **Fig. 1h** to **Fig. 1i**).

**Table 1.** Proteoforms identified from *scPiMS* and proteoform imaging of rat hippocampus.

| Protein name | Gene | UniProt<br>Accession | Average<br>mass | Modifications | E-value<br>(# of matched<br>fragments) |
| --- | --- | --- | --- | --- | --- |
| ATP synthase subunit epsilon, mitochondrial | ATP5F1E | P29418 | 5635.1 | Chain@2-51 | $2.3 \times 10^{-11}$ (8) |
| Polyubiquitin-B | UBB | P0CG51 | 8565.6 | Chain@1-76 | $1.90 \times 10^{-39}$ (37) |
| Acyl-CoA-binding protein | DBI | P11030 | 9938.0 | Chain@2-87,<br>Acetyl@N | $1.30 \times 10^{-15}$ (14) |
| Dynein light chain 2, cytoplasmic | DYNLL2 | Q78P75 | 10260.0 | Acetyl@N | $1.10 \times 10^{-13}$ (12) |
| 10 kDa heat shock protein, mitochondrial | HSPE1 | P26772 | 10812.8 | Chain@1-76,<br>Acetyl@N | $3.90 \times 10^{-12}$ (21) |
| Myelin basic protein | MBP | P02688-4 | 14216.2 | Chain@2-128,<br>Acetyl@N,<br>Phosphoryl@112 | $2.00 \times 10^{-11}$ (8) |
| Myelin basic protein | MBP | P02688-4 | 14246.2 | Chain@1-128,<br>Acetyl@N | $3.60 \times 10^3$ (3) |
| Myelin basic protein | MBP | P02688-4 | 14331.1 | Chain@1-128,<br>Acetyl@N,<br>Phosphoryl@8 | $5.40 \times 10^3$ (3) |
| **Galectin-2** | LGALS2 | Q9Z144 | 14648.3 | Chain@2-130,<br>Acetyl@N | $4.40 \times 10^3$ (3) |
| **Myelin basic protein** | MBP | P02688-5 | 17149.8 | Chain@2-158,<br>Acetyl@N | $1.50 \times 10^{-1}$ (8) |
| **Myelin basic protein** | MBP | P02688-2 | 18412.3 | Chain@2-169,<br>Acetyl@N | $3.5 \times 10^{-4}$ (12) |
| **Myelin basic protein** | MBP | P02688-2 | 18491.4 | Chain@1-169 | $2.90 \times 10^0$ (7) |
| **Brain acid soluble protein 1** | BASP1 | Q05175 | 21896.3 | Chain@2-220,<br>Acetyl@N,<br>Myristoyl@1) | NA (12) |
| **Superoxide dismutase [Mn], mitochondrial** | SODM | P07895 | 22291.1 | Chain@25-222 | NA (5) |
| **Malate dehydrogenase, mitochondrial** | MDHM | P04636 | 33182.1 | Chain@25-338 | NA (30) |
| **Fructose-bisphosphate aldolase A** | ALDOA | P05065 | 39219.8 | Chain@2-364 | NA (7) |
| **Alpha-enolase** | ENO1 | P04764 | 47040.2 | Chain@2-434 | NA (14) |
| **Gamma-enolase** | ENO2 | P07323 | 47051.2 | Chain@2-434 | NA (9) |

### scPiMS: insights into scProteoform heterogeneity

**Fig. 2a** shows the mass spectrum of the aggregated 10.8k cell dataset with some of the proteins annotated for which proteoforms were assigned and identified. Myelin basic proteins (MBPs) are among the most abundant in the dataset with a complex landscape of methylation, phosphorylation along with oxidations that can be introduced artefactually in cell processing. A typical MBP scProteoform detection is shown in **Fig. 2b** containing 66 matched ions and (PAS = 1.00). Aside from proteoforms involved in primary metabolism (glycolysis, the TCA cycle, ATP synthase and OXPHOS), we also detected a variety of brain-specific proteoforms including Alpha-endosulfine and Corticoliberin (**Fig. 2a**, Table 1). Moreover, we identified several proteoforms expressed by specific cell types in the rat hippocampus, in particular, neuronal (*e.g.*, Alpha-synuclein-SYUA, Ubiquitin carboxyl-terminal hydrolase isozyme L1-UCHL1, Gamma-enolase-ENOG) and astrocytic proteins (*e.g.*, Protein S100B, multiple isoforms of Glial fibrillary acidic protein, GFAP, Fructose-bisphosphate aldolase C-ALDOC). Example scProteoform detection of SYUA, ALDOC, and ENOG cell markers are shown in **Fig. 2b-e** with the detection of 2-66 single ions and PAS scores ranging from 0.5-1.0. This highlights the potential of *scPiMS* in high-throughput cell type classification without the need of antibodies or labels.

**Fig. 2.**
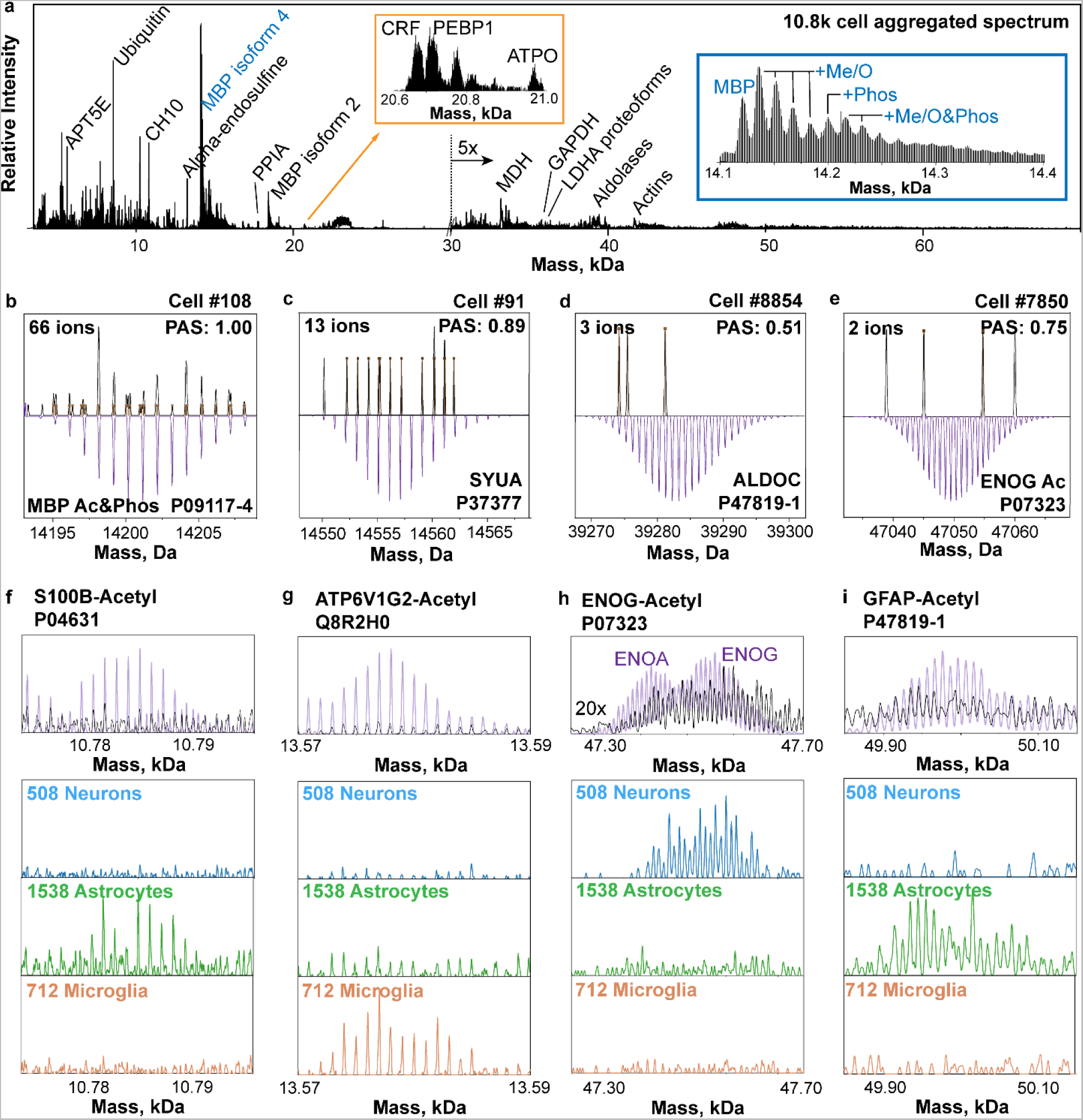
Mass spectra obtained from 10836 aggregated cells, single cells, and specific rat brain cell types. (a) aggregated 10.8k single cell spectrum in the 4-70 kDa mass range labeled with identified proteoforms. Insets show expanded regions around 21 kDa (orange box) and 14 kDa, with the latter capturing the proteoform landscape of Myelin basic proteins (blue box). (b-e) Single-cell mass spectra (dark trace) showing the detection of single ions (top) of four selected scProteoforms and their theoretical isotopic distributions (bottom, mirrored in purple). (f-i) a set of 4 proteoform markers detected in *scPiMS* that differentiate and provide initial assignment of rat hippocampal cell types (508 neurons, 1538 astrocytes and 712 microglia). In each panel, the top spectrum shows an expanded spectral region of the ion library (light purple) and 10.8k-cell aggregated dataset (black trace); the bottom three insets show proteoform spectral regions, all normalized to the same abundance scale, aggregated from 508 neurons, 1538 astrocytes and 712 microglia, respectively.

We sought next to differentiate major cell types from the individual 10.8k cells based on spectral features. To this end, a subset of the 13 identified protein markers reported to show differential expression in neurons, astrocytes, and microglia was assembled. Candidate cell marker proteoforms were validated against the aggregated single cell spectra, and proteoforms with relatively high abundance and little or no overlap were selected. We ranked the 10.8k cells according to a simple binary score for initial assignment of neurons, astrocytes, or microglia (Methods). From the highest scoring subpopulations, we assigned a putative subset of 508 neurons and 1538 astrocytes, and 712 microglia. Among the marker proteoforms detected in specific cell type populations, GFAP is a well-known immunohistochemistry marker for astrocytes.^26, 31^ The aggregated mass spectra from the three cell types show significant differential abundance in four examples of proteoform markers of cell type (**Fig. 2f-i**).

### Conclusion

A new style of single-cell proteomics was demonstrated on ∼10,000 cells with expediency. While conceptually similar to direct MS studies of single cells in the past, this work translates the potential of single ion technology into a scalable approach to single-cell proteomics for fast molecular profiling and identification with proteoform specificity up to ∼70 kDa. The new *scPiMS* approach also converts the ‘no-digestion’ or ‘top-down’ philosophy to proteomics into a key advantage that opens a major bottleneck in SCPs by >20-fold (∼2000 cells/day compared to ∼150 cells/day^24^), yet involves interpretation of a new data type in the field. This throughput will enable proteoform signatures from rare cells to be captured. As shown here, the *scPiMS* platform will be advantaged tremendously using a reference set of proteoforms and their ions (*c.f.*, inclusion into the Human Proteoform Atlas^32^) to both accelerate the proteoform annotation step and the depth the analysis obtained in each cell. By providing proteoform-level information, *scPiMS* promises to advance single cell biology by mapping cell types underlying diverse disease phenotypes. To the extent proteoform measurement connects to complex phenotypes, a mature version of *scPiMS* may create an outsized impact in single cell biology provided it can find wide adoption in a manner analogous to scRNA-seq.

## Materials and Methods

### Tissue Preparation

Three 2-month-old male Sprague Dawley outbred rats (*Rattus norvegicus*, Inotiv, West Lafayette, IN) were used in experiments. Animals were fed *ad libitum* and housed in a 12-h light cycle. Animals were asphyxiated using CO2. All procedures were performed in accordance with animal use protocol approved by the University of Illinois Institutional Animal Care and Use Committee, and in compliance with both federal and ARRIVE guidelines for the humane treatment of animals.

Immediately after euthanasia, animals were perfused transcardially with ice cold modified Gey’s balanced salt solution (mGBSS) containing (in mM): 1.5 CaCl2, 5 KCl, 0.2 KH2PO4, 11 MgCl2, 0.3 MgSO4, 138 NaCl, 28 NaHCO3, and 0.8 Na2HPO4, and 25 HEPES, pH 7.2. One third of the hippocampus from one hemisphere was surgically dissected for single cell preparation. Intact hemispheres of the same brains as used for cell population collection were frozen and sectioned for intact tissue PiMS imaging. Rat hippocampus tissue punches and thin sections were prepared according to published protocols.^33^ Rat brain hemisphere slices 16 µm thick were cut at -16°C using a cryostat (Leica CM3050 S, Leica Biosystems GmbH, Wetzlar, Germany). Brain slices containing hippocampal areas of all three animals were deposited on the same slide. Sections were rinsed with 200 Proof ethanol twice. Tissue samples were stored at −80°C before imaging analysis by *scPiMS* using the published approach.^21^

### Preparation of Individual Cell Populations

We used the procedures for single cell isolation developed for single cell metabolomics, adapted here for *scPiMS*.^26, 34^ One-third of the hippocampus from one hemisphere was surgically dissected and treated with the papain dissociation system (Worthington Biochemical, Lakewood, NJ) for 120 min. at 34°C with superficial oxygenation. After the treatment, tissues were gently rinsed twice with ice cold mGBSS. Mechanical tissue dissociation was performed in ice cold mGBSS supplemented with Hoechst 33342 (1 μg/mL). The individual cell suspension was deposited onto indium tin oxide (ITO) glass slides (Delta Technologies, Loveland, CO) to achieve low density cell population with individual cells spaced by an average of >200 um. Cell populations from three different animals were deposited onto separate marked areas on ITO glass slides. Cells were sedimented and adhered to slide surface for 10-20 min. Surrounding cell media was quickly replaced with mGBSS containing 33% glycerol. After brief incubation for ∼10-20 s most of the solution present on ITO glass slide was removed. Slides were rinsed twice with 200 Proof ethanol (Lab Alley, Spicewood, TX).

### Fluorescent Microscopy Experiments

Images of populations of single cells were taken in mixed bright-field and fluorescence mode using a Zeiss Axio M2 microscope (Zeiss GmbH, Jena, Germany) equipped with an AxioCam ICc5 camera, X-cite Series 120 Q mercury lamp (Lumen Dynamics, Mississauga, ON, Canada) and a HAL 100 halogen illuminator (Zeiss). The DAPI (ex. 335–383 nm; em. 420–470 nm) dichroic filter was used for fluorescence excitation. A 2.5× objective was used for fast acquisition with a 13% overlap between individual images allowing proper image stitching and formation of a single view for entire glass slide. Images were processed and exported as big .tiff files using ZEN software version 2 blue edition (Zeiss). Numbers of cells on slide in regions of interest were determined using ImageJ.^35^

### scPiMS probe fabrication and ion source conditions

A custom-designed nano-DESI source was used for all data acquisition. The experimental details of nano-DESI MSI have been described elsewhere.^36, 37^ Briefly, the nano-DESI probe is comprised of a flame-pulled fused silica primary (OD 40 µm, ID 20 µm, Molex, Thief River Falls, MN) and a nanospray capillary (OD 150 µm, ID 40 µm) with the spray side of the nanospray capillary positioned towards the MS inlet. The nano-DESI probe utilizes a dynamic liquid bridge formed between the capillary junction and the glass surface to extract analytes when brought into contact with the glass surface. The liquid bridge is dynamically maintained by solvent propulsion from the primary capillary and instantaneous vacuum aspiration through the nanospray capillary. All samples were electrosprayed in positive ion mode under denaturing conditions in a 60%/39.4% acetonitrile/water and 0.6% acetic acid solution compatible with both protein extraction and ionization. The solvent flow rate was kept in the range of 160-500 nL/min. The ion source conditions on the mass spectrometer were set as follows: ESI voltage: +1.7-3 kV; in-source CID: 15 eV; S-lens RF level: 80%; capillary temperature: 360°C.

### scPiMS operation modes

scPiMS experiments were performed in two distinct modes: targeted mode and high-throughput scanning mode. In targeted mode, a single cell feature was first found by the 900× high-magnification microscope (Supplementary Fig. 1). In particular, a nano-DESI probe was brought into contact with the glass surface at a location ∼50 µm away from the single cell to form the liquid bridge, and was subsequently moved toward the cell with manually-controlled stage motions after MS data acquisition was triggered. The probe was typically parked on the cell for one minute to obtain a targeted “cellogram” (chronogram of total ion counts). The cellogram typically starts from a drastic rise in protein signal followed by an exponential decay over time, which is characteristic of the extraction kinetics of cellular protein analyzed by *scPiMS*.

High-throughput scanning mode was performed by moving the glass slide under the nano-DESI probe in parallel lines at a constant linear velocity. The spacing between adjacent lines was set similar to the size of the liquid bridge formed by the probe and the surface to ensure all the surface area containing single cells were covered. The size of the liquid bridge in nano-DESI can be adjusted in range from 25 to 400 µm depending on the size of the capillaries used to fabricate the probe. For *scPiMS*, 200 µm was selected considering the throughput of the experiment to covering the entire area containing single cells. Larger probes will result in multiple adjacent cells on the surface being analyzed at the same time, which obscures the protein profiles from these individual single cells. Statistical modeling was carried out to simulate the percentage of single cell extraction events using the single cell coordinates obtained from fluorescence microscopy. In particular, surface area containing single cells was divided into 200 µm square grids, and number of grids that contain one cell or more than one cell were calculated. At the density of cells loaded on the glass slides, 90% of the cells can be sampled as single cell features using the 200 µm probe (**Fig. 1f**).

### Cellogram optimization

A cellogram obtained in targeted mode revealed the extraction kinetics of *scPiMS* probe on single cells and provided guidance to optimizing the parameters in high-throughput scanning mode. In particular, a targeted mode cellogram was obtained by parking the probe at the cell and monitoring the total ion response. The time at which the targeted cellogram falls back to the baseline indicated that readily extractable proteins were captured. This gave the approximate time that the cell should be exposed to the probe and informs the overall throughput for *scPiMS*. More importantly, in high throughput scanning experiments, extraction of protein content from a single cell before getting in contact with the next cell guarantees that the signal carryover between adjacent cells is minimized. The upper limit was employed to calculate the “exposure time” of a single cell to the liquid bridge in the rastering mode. The average exposure time for single cells at a 500 nL/min. solvent flow rate was found to be 5 s from a few targeted cellograms. The rastering scan rate was determined by the size of the liquid bridge, size of the single cell, and the exposure time (Eq. 1). Considering the small size of the rat brain single cells (∼10 µm) compared to the liquid bridge (200 µm) in the current *scPiMS* configuration, the size of the liquid bridge was used to calculate the rastering scan rate (Eq. 1):

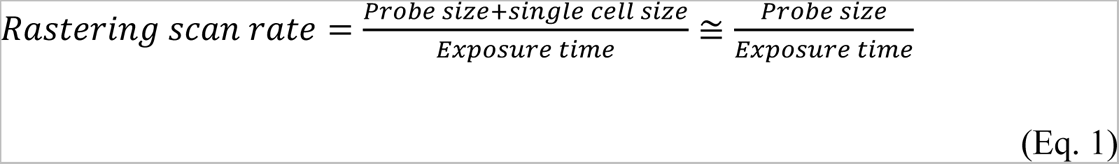

A 40 µm/s scan rate (200 µm divided by 5 s) was used for all the 10.8k *scPiMS* runs. Interestingly, the exposure time was found to be flow-rate dependent. In particular, in precursor 2k single cell runs, a reduced solvent flow rate at 180 nL/min was employed, which gave an average exposure time of 8 s. We reasoned that the solvent replenishing rate and band diffusion in the liquid flow in the secondary capillary both contribute to the observed slower kinetics of extraction. This further suggests that targeted cellograms should be obtained before starting the high-throughput scanning experiments if *scPiMS* sampling conditions were changed.

### scPiMS Data Acquisition

*scPiMS* data acquisition was performed in the individual ion mass spectrometry (I^2^MS) mode on a previously described Orbitrap Q-Exactive Plus mass spectrometer (Thermo Fisher Scientific GmbH, Bremen, Germany).^24^ The spectral acquisition rate was set at one scan per second. During *scPiMS* data acquisition, proteoforms from single cells were sampled and ionized by a nano-DESI probe to generate multiply-charged ions distributed across multiple charge states. The ion injection time was optimized such that ions in one detection period were collected in the individual ion regime, which gives a singular ion signal at a defined *m/z* (or frequency) value. Due to the protein signals emerging from single cells, the majority of the *scPiMS* ion detection periods were dominated by individual ion signals even at extended MS injection times over 200 ms.

As discussed in previous work, the HCD pressure level has a strong impact on the ion accumulation and survival performance in I^2^MS. The HCD pressure level was typically set at 0.5 (UHV pressure <5 × 10^-11^ Torr) to reduce collision-induced ion decay within the Orbitrap analyzer without sacrificing the trapping efficiency for intact protein ions. The HCD pressure setting of 0.2 was used throughout the *scPiMS* experiments to reduce the chemical noise collected in the spectrum at extended MS injection times.

In addition, the Orbitrap central electrode voltage was adjusted to -1 kV to improve the ion survival rate in I^2^MS. Additional relevant data acquisition parameters were as follows: mass range: 500-1500 *m/z*; AGC mode: disabled; enhanced Fourier transform: off; averaging: 0; microscans: 1. Time-domain data files were acquired at detected ion frequencies and recorded as Selective Temporal Overview of Resonant Ions (STORI) files.^38^ STORI setting: Enabled.

### scPiMS feature selection & data analysis

*scPiMS* data analysis was performed using MATLAB and C^#^ script developed in-house. *scPiMS* raw data were collected as continuous chronograms containing discrete or overlapping “cellograms” and other peak features from chemical noise on the glass slide collected during the experiments. Cellogram picking was first performed to recognize MS scans corresponding to true single cell events and eliminate the blank and noise scans from the dataset. We first took advantage of STORI analysis in I^2^MS processing to stratify the “protein-signal-like” and “noise-like” scans. In particular, typical chemical-noise-like ions from the ion source show lower ion lifetime and lower R^2^ in their STORI plots. We found the R^2^ cutoff of 0.999 as a suitable threshold to achieve sufficient signal-to-noise ratio for extraction of true single cellograms.

The ion chronogram constructed at R^2^ >0.999 was employed for peak picking with a peak intensity threshold of 10 and a minimum peak distance of 5 (exposure time * MS scan rate). In specific cases, the threshold was set to 50 and the R^2^ cutoff was set to 0.9996 to take into account the variation of the noise level in the dataset on different days of acquisition when instrument performance changed notably. The peak features picked by the program were further categorized into single-cell multi-cell and cellogram-like features using peak characteristics. In particular, if the half-width-at-half-maximum on the right side of a cellogram was more than two scans (2 s), the feature was recognized as a multi-cell feature. Multi-cell features were strongly filtered in this step and only the single cell features were used for further analysis. The number of single cell features was also validated and matched to the fluorescent single cell feature count as described in the “*scPiMS operation modes*” section. We observed in rare cases that the single cell feature count is substantially lower than the simulated fluorescent count. This was attributed to aberrant probe conditions, and the peak picking criteria were not further adjusted to match the fluorescent features. From a total of 15.1k single cell features, we were able to detect 10.8k single-cell-features over the course of 10 days.

The STORI files of the MS scans from the single-cell-like features were reextracted from the dataset and concatenated to construct a mass-domain spectrum. In particular, all individual ion signals were resubjected to charge state assignment for mass calculation. The neutral masses of the protein ions were calculated by:

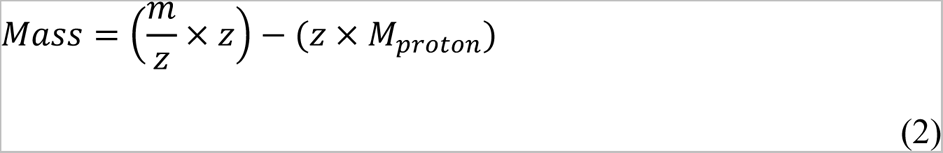

Charge state (*z*) is obtained from the slope of induced image current determined by the STORI analysis described in detail elsewhere.^38^ Accurate charge assignment of each ion was statistically evaluated by comparing the slopes of its isotopologues across different charge states from the entire dataset. We employed a probability metric to filter out ions with a lower probability score and construct mass-domain isotopic distribution with statistical confidence.

### scProteoform scoring and false discovery rate (FDR)

Individual ions corresponding to each single cell feature were aggregated into a single cell spectrum and were utilized to generate a score for each proteoform in a given cell feature. The proteoform assignment space may be determined via multiple pathways; either isotopic envelopes created via the THRASH deconvolution algorithm^39^ or envelopes created and validated through identified proteoforms from intact mass tag (IMT) search or MS/MS via the PiMS platform described in detail in the “*Intact Mass Tag (IMT) Search & Gene Ontology (GO) analysis”* and *“Data analysis and Proteoform MS/MS along with Identification*” section. Cell specific individual ions were searched against the isotopic envelope library and matched to specific isotopic peaks with a tolerance of ±0.3 Da, these ion matches are scored and combined with all ion scores for a given proteoform to yield a single cell proteoform score (PAS) using the following equation:

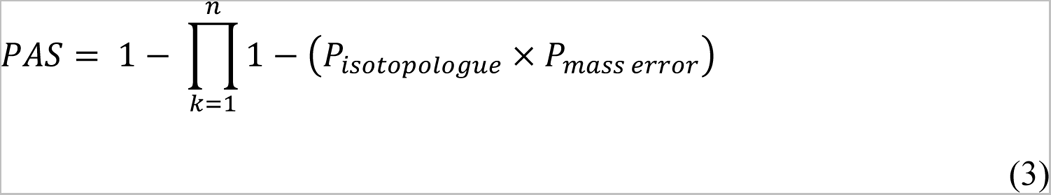

Where *P*_*isotopologue*_ is the expected relative intensity of the matching isotopologue and *P*_*mass error*_ is a probability score for the mass error between the observed individual ion mass and the theoretical mass of the isotopologue, utilizing the cumulative distribution function (CDF) of a normal distribution with a mean and sigma determined by the theoretical mass and width of a isotopic peak respectively:

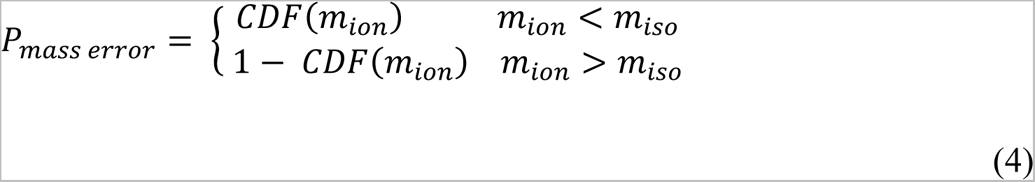

Where *m*_*ion*_ is the mass of the individual ion and *m*_*iso*_ is the theoretical mass of the matched isotopologue. To perform multiple hypothesis testing, an empirical false discovery rate (FDR) procedure was implemented at the single cell level. Decoy proteoforms (10 per proteoform in the library) were generated using random amino acids and constrained to be the same sequence length as the representative proteoform. These decoys were scored alongside the proteoform hits, rank ordered and all proteoform hits given a *q*-value. Proteoforms with a *q*-value less than 0.05 (5% FDR) were classified as assigned in that single cell.

### Cell Type Marker Scores

Using the FDR controlled proteoforms scores for various cell type markers for neurons, astrocytes and microglia, we were able to create a cell type score for each cell. Marker proteoform scores associated with a given cell type were combined with the following:

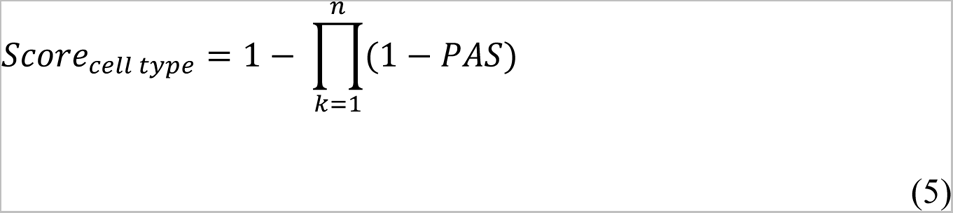

Where *n* is the number of markers used for the particular cell type, PAS is the proteoform assignment score. For each cell, a cell type score is generated for each of the three cell types being probed. Cells were assigned a cell type when they met the following criteria:

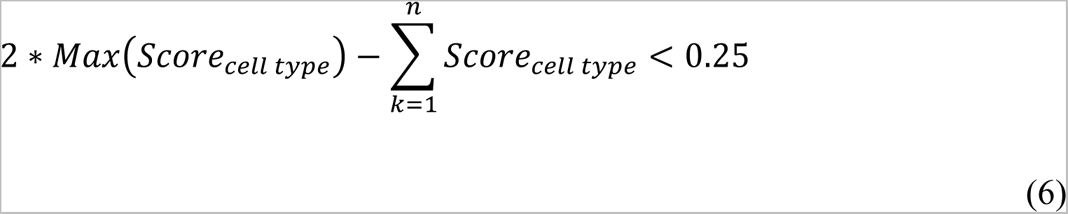

### Intact Mass Tag (IMT) Search & Gene Ontology (GO) analysis

The summed mass-domain ion library was converted to *.mzML* format and processed using a custom version of TD Validator (Proteinaceous, Evanston, IL) implemented with an MS^1^ IMT search function. The library spectrum was self-calibrated by +10.25 ppm according to the accurate masses of six identified proteoforms in the 10-50 kDa mass range. A custom protein database was constructed from top 1000 most abundant proteins in a bottom-up proteomics study of mouse brain hippocampus was used for the search.^40^ In particular, the protein names of the top 1000 mouse proteins were transformed into 764 corresponding rat protein entries. The IMT search was performed with a ±1.5 ppm mass tolerance considering Methionine on/off, monoacetylation, and monophosphorylation as possible proteoform modifications in the database (*i.e.*, a search space of ∼5000 proteoforms). Additional proteoform matches were curated by spectral inspection and manual annotation of putative modifications recorded in the hippocampal-specific top-down proteomics study by Fournier *et al*.^28^ and the SwissProt rat proteome database, resulting in a total of 169 proteoform matches. Gene Ontology (GO) analysis using Metascape (https://metascape.org/) was performed by retrieving a list of Entrez Gene ID for the 169 identified proteoforms using the ID mapping tool on UniProt.^41^

### Data analysis for Proteoform Identification by MS/MS along with Identification

Targeted MS/MS experiments were performed in the hippocampal region of a brain tissue section from one of the animals where single cells were obtained. Briefly, a survey PiMS line scan along the hippocampal region was obtained and processed using I^2^MS that gives the *m/z*, charge state and favorable location for MS/MS experiment for a list of proteoform targets.

MS/MS experiments were performed on an adjacent line region offset by 50 µm according to a multiplexed acquisition method generated from the last step. A 0.8 *m/z* isolation window was employed for most of the targets. The raster scan rate for survey and MS/MS experiments was 3 µm/s. MS/MS data acquisition was conducted using higher-energy collisional dissociation (HCD) with fragment ion detection in either traditional ensemble or I^2^MS mode for <20 kDa and >20 kDa targets, respectively. Ensemble MS/MS experiments were performed on an Orbitrap Exploris 480 mass spectrometer (Thermo Fisher Scientific, Bremen, Germany) using a mass resolution of 30000 at an acquisition rate of 14.5 spectra/s and a HCD collision energy of 25-45 eV. MS/MS in I^2^MS mode were performed on a Q Exactive Plus described in the “*scPiMS Data Acquisition*” section using an Orbitrap detection period of 1 s (HCD pressure setting = 0.5).^42^ Typical values for collision energy and injection time used in this study were 12 eV and 700 ms, respectively.

Total data acquisition time for each target ranged from one to five minutes. For database search of targets collected in I^2^MS mode, mass-domain spectra were generated for database search. Top-down MS/MS searches were performed using ProSight Native and TDValidator (Proteinaceous) against a set of proteoforms created from the entire rat SwissProt database. Proteoform identifications were reported using E-values and number of matched fragments for ensemble and I^2^MS data type, respectively.

---

**Supplementary Fig. 1.**
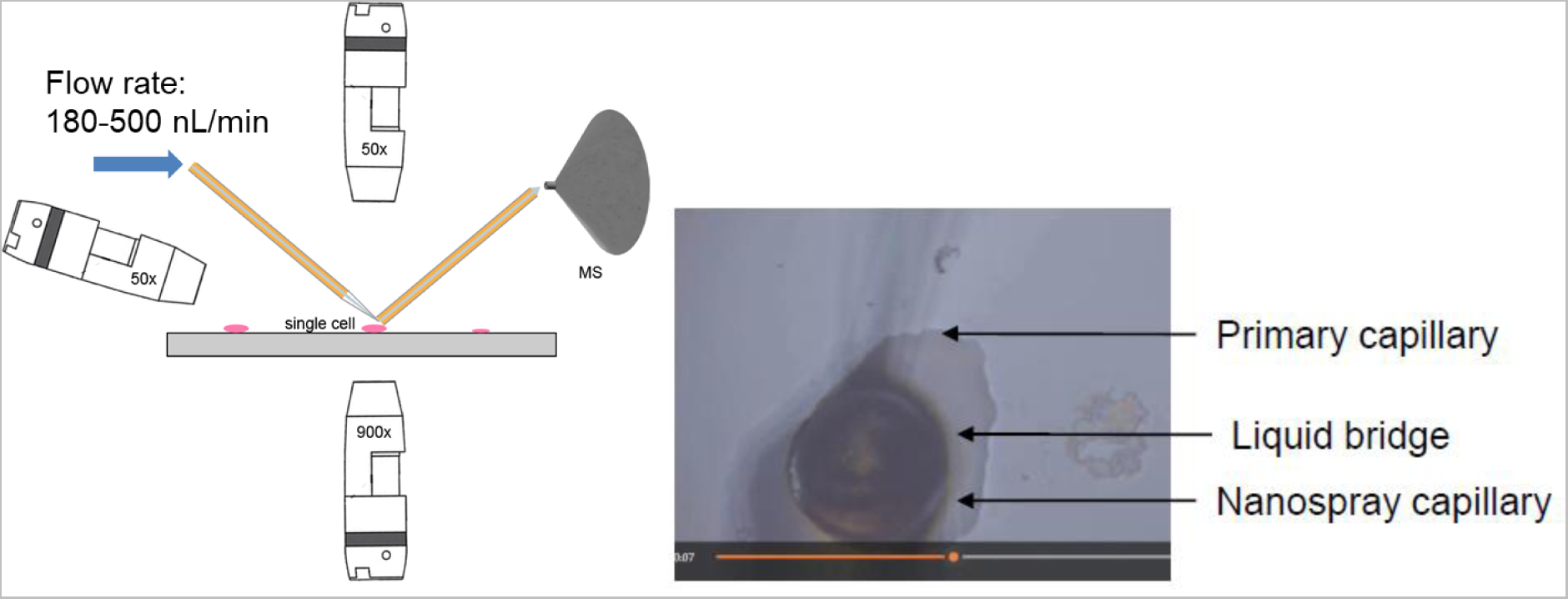
Experimental setup for *scPiMS* in targeted mode.

## Funding

National Institutes of Health P41 GM108569 (NLK)

National Institutes of Health UH3 CA246635 (NLK)

National Institutes of Health P30 DA018310 (NLK and JVS)

National Institutes of Health P30 CA060553 (awarded to the Robert H. Lurie Comprehensive Cancer Center)

## Author contributions

Conceptualization: NLK

Methodology: PS, NLK, JOK

Resources: NLK

Software: PS, MARH, JBG, BPE, RTF

Investigation: PS

Visualization: PS, MARH, JBG, BPE, JOK

Supervision: JOK, NLK

Writing—original draft: NLK

Writing—review & editing: PS, MARH, FAB, JBG, BPE, RTF, JOK, NLK

## Data and materials availability

Custom compiled code used to process and create I^2^MS files is available.^43^ Additional software and data that support the findings of this study are available from the corresponding author upon request.

